# Physiological variation reflects bioclimatic differences in the *Drosophila americana* species complex

**DOI:** 10.1101/453571

**Authors:** Jeremy S. Davis, Leonie C. Moyle

## Abstract

**Background:** Disentangling the selective factors shaping adaptive trait variation is an important but challenging task. Many studies—especially in *Drosophila*—have documented trait variation along latitudinal or altitudinal clines, but frequently lack resolution about specific environmental gradients that could be causal selective agents, and often do not investigate covariation between traits simultaneously. Here we examined variation in multiple macroecological factors across geographic space and their associations with variation in three physiological traits (desiccation resistance, UV resistance, and pigmentation) at both population and species scales, to address the role of abiotic environment in shaping trait variation.

**Results:** Using environmental data from collection locations of three North American *Drosophila* species—*D. americana americana, D. americana texana* and *D. novamexicana*—we identified two primary axes of macroecological variation; these differentiated species habitats and were strongly loaded for precipitation and moisture variables. In nine focal populations (three per species) assayed for each trait, we detected significant species-level variation for both desiccation resistance and pigmentation, but not for UV resistance. Species-level trait variation was consistent with differential natural selection imposed by variation in habitat water availability, although patterns of variation differed between desiccation resistance and pigmentation, and we found little evidence for pleiotropy between traits.

**Conclusions:** Our multi-faceted approach enabled us to identify potential agents of natural selection and examine how they might influence the evolution of multiple traits at different evolutionary scales. Our findings highlight that environmental factors influence functional trait variation in ways that can be complex, and point to the importance of studies that examine these relationships at both population- and species-levels.

## Background

Determining how environmental variation shapes trait variation within and between species is central to our understanding of how natural selection can drive adaptive change. One hallmark of adaptation is a consistent association between trait variation and one or more aspects of the natural environment. Classically, these associations have been assessed via the study of clines; by definition, clines exhibit spatial variation, and geographic space is frequently environmentally heterogeneous, so traits exhibiting functionally-relevant clinal variation are clear candidates for targets of local selection. Latitudinal or altitudinal clines have received particular attention in numerous systems, including *Drosophila, Arabidopsis thaliana*, and humans, where analyses indicate strong trait-environment associations for several physiological and other variants (Adrion et al. 2015, Fournier-Level et al. 2011, Hancock et al. 2011). Nonetheless, even among these well-characterized examples, the underlying cause of clinal variation is still not always clear, particularly when trait variation is surveyed across generalized geographic space as opposed to specific environmental gradients.

Macroecological analyses are one useful method for connecting environmental variation to trait adaptation. Using environmental data from GIS-based databases, these approaches quantify the direction and magnitude of bioclimatic variation across species ranges. Investigating how these macroecological factors co-vary with trait variation between populations can identify which aspects of the environment might be most important for shaping population-level variation and provide insight into patterns of local adaptation (Kozak et al. 2008). Extending these analyses to include populations from multiple species across space allows further investigation into how environment is influencing the evolution of phenotypic differences that manifest at both the species and population levels.

In *Drosophila,* patterns of intraspecific clinal variation and species differences have pointed to several traits as potential targets of environmentally-mediated selection (Adrion et al. 2015). A long history of latitudinal analyses in North American and Australian *Drosophila melanogaster* have revealed clinal variation in body, egg, and wing size, bristle size, ovariole number, lifetime fecundity, cold tolerance, and diapause incidence among other traits (Zwaan et al 2000, Coyne and Beecham 1987, Azevedo et al. 2002, 1996, David and Bocquet 1975, Schmidt et al. 2005, Schmidt and Paaby 2008, reviewed Adrion et al. 2015). More broadly across *Drosophila*, multiple studies have shown an association between pigmentation variation and latitude, including within *D. melanogaster* in Europe (David et al. 1985), Australia (Telonis-Scott et al. 2011), India (Munjal et al. 1997), and sub-saharan Africa (Pool and Aquadro 2007, Bastide et al. 2014), as well as within *D. simulans* (Capy et al. 1988) and the *D. cardini* group (Heed and Krishnamurthy 1959). However, despite this wealth of data, in many cases the environmental and selective factors responsible for driving clinal variation in these traits are equivocal, and sometimes conflicting. For example, latitudinal studies on thoracic pigmentation in *D. melanogaster* have implicated seasonal and annual temperature variation as the primary selective agent explaining positive correlations with latitude in Europe (David et al. 1985), Australia (Telonis-Scott et al. 2011), and India (Munjal et al. 1997)—although these patterns can covary with other factors such as altitude (e.g. in Africa; Pool and Aquadro 2007)—while clinal UV intensity variation has been invoked to explain an opposing pattern seen in Africa (Bastide et al. 2014). These abiotic factors are proposed to shape traits directly via selection to increase physiological resilience where environmental conditions are most stressful. In addition, these factors have also been proposed to shape the relationship between traits, due to potential pleiotropic effects that changes in cuticle structure might have on multiple physiological stress responses, including both UV and desiccation resistance, as well as on pigmentation. For instance, patterns of pigmentation variation have frequently been proposed to be explained by associated desiccation resistance variation, with studies showing that increased desiccation resistance is correlated clinally with darker pigmentation in *D. polymorpha* (Brisson et al. 2005), *D. ananassae* (Parkash et al. 2010), and Indian *D. melanogaster* (Parkash et al. 2008), although this pattern was not observed in *D. americana* (Wittkopp et al. 2009). Matute and Harris (2013) found no relationship between desiccation resistance and pigmentation in *D. yakuba* and *D. santomea* but observed that lighter pigmentation confers greater UV resistance—a result that runs contrary to implied latitudinal patterns in other species. Accordingly, despite the attention received by these traits, their relationships to potential environmental agents and to each other remains poorly understood in many species.

The *Drosophila americana* group provides a good system for investigating how environmental variation across large spatial regions could influence physiological adaptation within and between species. This group consists of three members of the *virilis* clade native to North America—the two subspecies *Drosophila americana americana* and *Drosophila americana texana*, and their sister species *Drosophila novamexicana. D. novamexicana* is localized to the arid southwestern US, while *D. a. americana* and *D. a. texana* each span a wide geographic and climatic range from the great plains in the west, across to the east coast of North America (Figure 1). While *D. novamexicana* is clearly spatially differentiated from the two subspecies of *D. americana*, in the absence of quantitative data it’s unclear which of many covarying factors might represent the strongest differences in habitat between this species and its relatives. Similarly, while the *D. americana* subspecies are generally distributed on a north (*D. a. americana*) to south (*D. a. texana*) cline, their ranges show substantial overlap (McAllister 2002), and the magnitude and nature of their climatic differences have not previously been quantified. Further, these species show evidence for variation in both pigmentation and desiccation resistance traits (Wittkopp et al. 2011, Clusella-Trullas and Terblanche 2011), but the relationship between this variation and macroecological factors within and between species remains unclear.

**Figure 1:**
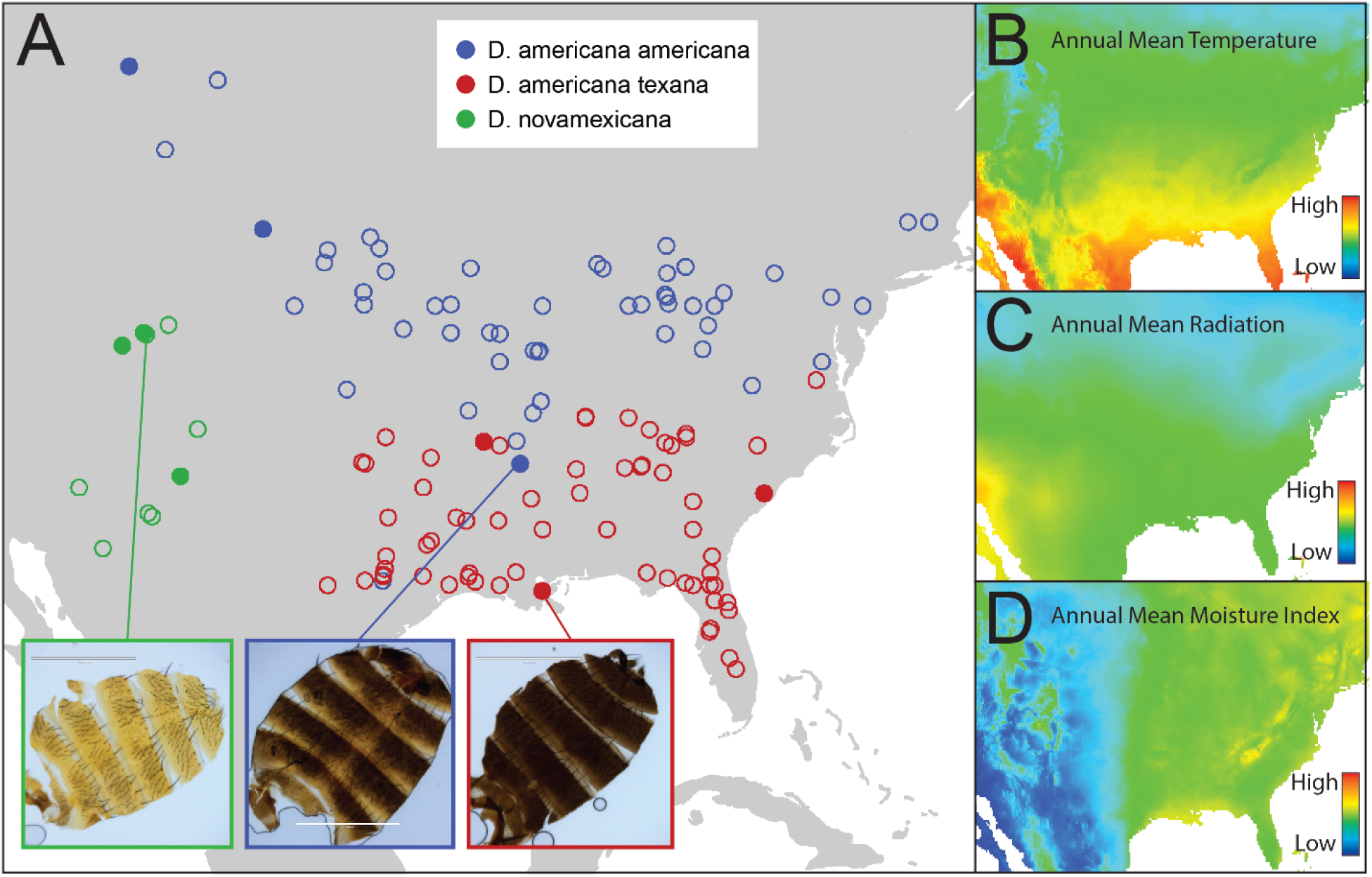
Distribution of collection locations and major environmental variables for our three focal species. Panel A shows collection records for each of *D. novamexicana* (green), *D. a. americana* (blue), and *D. a. texana* (red) as obtained from the TaxoDros database (taxodros.uzh.ch, see methods). Closed circles indicate the nine sample locations for populations used in this study. Panel B-D show heatmaps of spatial variation in annual mean temperature (B), annual mean radiation (C), and annual mean moisture index (D), as obtained from the Worldclim and Climond databases. Cuticle dissection images inset into Panel A are representative male cuticles from (left to right) Grand Junction, Colorado, White River, Arkansas, and New Orleans, Louisiana.

Here, our goal was to assess whether broad bioclimatic factors shape physiological variation in the *D. americana* species complex. To do so, we quantified the major axes of environmental variation within and between species, using climate variables from known occurrence locations. Based on these major axes, we assessed variation in three relevant physiological traits—desiccation resistance, UV resistance, and pigmentation—in a nine focal populations, to investigate evidence for associations between trait variation and this putative selective environmental variation. We found evidence for differentiation among these species in two primary axes of environmental variation, as well as for desiccation resistance and pigmentation, although patterns of association differed between these traits. We infer that species-level trait variation is consistent with natural selection imposed by habitat differences—in particular the influence of moisture availability on variation in desiccation resistance.

### Methods

#### Experimental Fly stocks

Three stocks from each focal species were obtained from the University of California San Diego Drosophila Species Stock Center (DSSC). We used *Drosophila novamexicana* stocks from San Antonio, NM, Grand Junction, CO, and Moab, UT (15010-1031.08, 15010-1031.00, and 15010-1031.04 respectively); *D. americana americana* (hereafter *D. a. americana*) stocks from Chinook, MT, Chadron, NE, and White River, AR (15010-0951.02, 15010-0951.06, and 15010-0951.17 respectively); and, *D. americana texana* (hereafter *D. a. texana*) stocks from New Orleans, LA, Jamestown, SC, and Morrilton, AR (15010-1041.24, 15010-1041.29, and 15010-1041.23 respectively). All stocks were collected between 1946 and 1950. *D. americana* is divided into subspecies according to presence of a chromosomal fusion of the X- and 4-chromosomes in *D. a. americana* that shows a distinct latitudinal cline (McAllister 2002). For simplicity we refer to them by their subspecies names. All fly stocks were reared on standard cornmeal media prepared by the Bloomington Drosophila Stock Center (BDSC) at Indiana University, and were kept at room temperature (∼22C).

#### Environmental data

To quantify environmental variation in the natural range of our three focal species, we extracted bioclimatic variable data from documented collection locations of each species and used these data to create principal components that summarize the major axes of climate variation. First, geographical coordinate data were obtained for all known collections using TaxoDros (www.taxodros.uzh.ch) – a database that compiles latitude and longitude coordinates from published collection records. After curating for duplicates and erroneous or unspecified coordinates, we retained passport data for 10 *D. novamexicana*, 73 *D. a. americana*, and 68 *D. a. texana* population locations. For each of these geographic locations, we extracted bioclimate variable data from two sources. From the Worldclim 2 database we extracted source location data at 30 arcsecond resolution for 19 bioclimatic variables (Fick and Hijmans 2017); and, from the CliMond archive (Kriticos et al. 2012) we extracted 16 additional bioclimatic variables at 10-minute resolution (see supplement). The latter were included despite this relatively coarse resolution because they contained additional data on ultraviolet (UV) radiation and soil moisture that were not available in the Worldclim registry. Many of these 35 bioclimatic variables describe alternative aspects of temperature, precipitation, and seasonality over different time intervals, so we performed a principal component analysis (PCA) in *R* on all 35 variables across all 149 population locations to reduce non-independence and redundancy in this dataset. PCA uses orthogonal transformation to generate a set of linearly uncorrelated principal components that summarize major axes of variation across a dataset of potentially correlated variables. Because the first 3 PCs explained ∼85% of the variation across populations (see Results), we used values for these PCs in our subsequent analyses of the relationship between environmental variation and variation in physiological traits.

#### Desiccation resistance assay

To assess population, species, and sex specific differences, desiccation resistance was assayed in replicate for individual males and females of each of our nine focal populations, using custom desiccation vials. Virgin flies were isolated within 24 hours of eclosion and aged individually for 3 days prior to the start of the experiment. Flies were then mouth aspirated individually into a modified *Drosophila* culture vial which contained a layer of 20g of Drierite, 10g of desiccated cork, and a piece of cheesecloth, and was sealed by a layer of parafilm. Each fly was placed above the cheese cloth and cork (in order to avoid direct contact with Drierite that negatively effects survival) and observed every 15 minutes until death. Death was assayed by observing the target fly for a total of 2 minutes, gently tapping the vial and watching for movement; when no limb or mouth movement occurred over that time, the fly was considered dead. Desiccation resistance was then quantified as the total time in minutes that each individual survived in a desiccation vial. A minimum of 5 replicates were performed per sex for each population. Trials were performed in blocks in which one fly of every identity (population x sex) was assayed simultaneously, to avoid confounding sex or population effects with trial date. At the end of the survival assay, each individual was weighed to obtain their dry weight (as a proxy for size) to include as a covariate in survival analyses. Dry (post-death) weight was determined to be an effective proxy for wet (pre-desiccation) weight in a pilot experiment in which individuals of each population and sex were weighed before and after individual desiccation trials (Pearson’s correlation, females: r(8) = 0.899, *P* < 0.001; males: r(8) = 0.925, *P* < 0.001).

#### UV irradiation resistance assay

We assessed UV-B resistance for each sex within each population (including the *D. virilis* line), at each of four different exposure intensities: 100, 500, 1000, and 5000 Joules/m^2,^ plus a control assay at 0 J/m^2^. UV resistance trials were performed similarly to Matute and Harris (2013) and Aguilar-Fuentes et al. (2008). Briefly, virgin males and females of each population were isolated and kept in single-sex groups of 20 for 24 hours prior to experiment start. Each group of 20 flies was then lightly anesthetized on a CO2 fly pad and weighed as a group before being irradiated with UV-B light at one of the four experimental intensities using an ultraviolet Stratalinker 2000 (Stratagene, La Jolla, CA). For the 0J exposure - which essentially measures longevity in the absence of acute UV exposure - flies were simply anesthetized, weighed, and placed in the Stratalinker without UV exposure. Each group was then transferred to a vial containing standard cornmeal media and scored once daily for number of flies still alive. Groups were transferred to fresh food vials as often as necessary—usually every seven days. The experiment continued until all flies in each vial were dead. Death was assessed here as in desiccation resistance assay above. For each assayed energy level, trials for both sexes in all ten lines were initiated simultaneously, to avoid confounding these factors with date effects.

#### Pigmentation assay

Dorsal abdominal pigmentation was assessed on individual males and females from each focal population in a similar manner to Wittkopp et al. (2011, dataset ‘A’). Briefly, individual 7-day old virgin flies for each sex and population were placed in 10:1 ethanol to glycerol mixture and stored at room temperature for 1-5 days. The dorsal abdominal cuticle (specifically tergites A3-A5) was dissected from each fly, isolated from other tissues, and mounted in Hoyer’s solution. Each cuticle was then viewed and digitally imaged on an AMG EVOS FL scope (AMG, Bothell, WA, USA) under standardized light conditions. Body color was quantified on gray-scale images of each cuticle by calculating the average median pixel intensity of 20 randomly-selected, non-overlapping regions on a 0-255 scale (avoiding the dorsal midline which has consistently lighter pigmentation), in Image J (NIH, Bethesda, MD, USA). Five replicate individuals from each sex within each population were assessed.

#### Statistical analyses

##### Environmental differences between species and populations

All statistical analyses were performed in R version 3.4.3, as was figure construction. We tested for evidence that species significantly differed in environment from one another by performing one-way ANOVAs with species as the independent variable and each of the first three PC axes as the dependent variables. These analyses were performed both on data from all collection localities used to generate the PC axes (N=149), and also with only the set of nine focal populations used for our trait analyses. For each analysis with a significant species effect, we also performed Tukey post-hoc contrasts to determine which species differed from one another.

##### Trait differences between sex, species, and populations

To assess the distribution of variation in our traits, we analyzed each physiological trait for differences between species, populations, and sex. For each of desiccation resistance and pigmentation, we fit a multi-way ANOVA with sex, species, and population nested within species, as independent variables, and each trait as the response variable. For desiccation resistance, dry weight was also included as a fixed effect to account for individual body size. For both of these traits we also performed post-hoc contrasts between each pair of species, using the Tukey test.

For UV resistance, effects of sex, species, and treatment level, were assessed using relative survival analysis and the R package relsurv (Pohar and Stare 2006). For each sex and population identity we used the 0J UV treatment exposure as the control (baseline) survival in a relative survival model, where relative survival in days following UV exposure is the response variable. We fit an additive model with sex, species, and treatment (energy level), as independent variables, to assess their contributions to UV resistance variation. Both species and treatment required a reference to be used, and we chose *D. americana* and 100J respectively as reference levels. (Results were unaffected by the specific choice of reference species, and only affected if 5000J was used as treatment reference.) Because we had only one trial per energy level for each sex within each population, we used our three populations per species to assess the effects of species identity on UV resistance. Finally, we used the median of a Meier-Kaplan curve estimate (Therneau 2013)—equivalent to the day in which 50% of the flies in a given trial are dead—as a summary statistic for UV resistance for each sex in each population at each given treatment level. These median values were used in subsequent analyses of trait-trait associations and trait associations with environmental PC axes (see below).

##### Environmental variation and association with physiological traits

We first examined how macroecological environmental variation (principal component axes) were related to desiccation resistance, UV resistance, or pigmentation variation across our nine focal populations, regardless of species. To do this we calculated Pearson’s correlation coefficient with mean population desiccation resistance survival time, UV resistance at each energy level, and pigmentation intensity as the response variable to either PC1 or PC2 values for our experimental populations. Then, because we observed that our PC axes exhibit statistical separation between species—that is, *D. novamexicana* had complete separation from the other two taxa along both PC1 and PC2 axes—we used a set of modified ANOVAs to evaluate how species and population identity influences trait-environment associations, for each sex separately. To do so, for each PC we first calculated the residuals from a one-way ANOVA with species as the independent variable, and then used these residual PC values in our analyses of population-level effects on each of our three traits. That is, for each of the first three PCs separately, we fit an ANOVA with residual PC values and species as independent variables, and the mean population trait value for either desiccation resistance or pigmentation as the response variable; a similar set of models were performed with UV resistance data from each UV treatment level, but median Meier-Kaplan curve estimates as the response variable. These analyses allowed us to simultaneously evaluate the contribution of both species differences and local environmental variation to variation in each physiological trait, and therefore assess how each PC contributes to variation in a given trait within each species. Because we performed 14 total tests, the Bonferroni-corrected significance level is p=0.004 for each trait.

Finally, we examined the strength of pairwise associations between each of our phenotypic traits of interest (desiccation resistance, pigmentation, and UV resistance at each of five levels), using Pearson’s correlation coefficients. Analyses were performed using population means (because each trait was measured on different individuals and, for UV resistance, groups of individuals), and on each sex separately (as there was evidence that each trait is moderately to strongly different between sexes; see results).

## Results

### Species differed along major axes of environmental variation

Our final dataset consisted of 149 latitude and longitude collection records across the United States (10 *D. novamexicana,* 71 *D. a. americana,* and 68 *D. a. texana*) for which we extracted 35 bioclimatic variables to generate principal component (PC) axes of environmental variation. These first 3 PCs explained 85% of the environmental variation across populations. Across all collection records (N = 149), we found that species differ along the environmental axes PC1 (*F*(2, 82.2), *P* < 0.001) and PC2 (*F*(2, 59.76), *P* < 0.001), but only marginally for PC3 (*F*(2, 3.4), *P* = 0.065). Post-hoc Tukey tests indicated that all 3 species differed from one another for both PC1 and PC2 (all *P* values < 0.001) (Figure 2). Despite modest power, species comparisons using environmental data from only our nine focal populations were similar: species differed along PC1 (*F*(2, 6.1), *P* = 0.0355) and PC2 (*F*(2, 5.6), *P* = 0.042), but not PC3 (*F*(2, 0.6), *P* = 0.564). For PC1, *D. novamexicana* focal populations differed from both *D. a. americana* (*P* = 0.033) and *D. a. texana* (*P* = 0.003), although the two *americana* subspecies populations did not differ (*P* = 0.070); for PC2, *D. novamexicana* differed from *D. a. americana* (*P* = 0.045), but the other two contrasts (*D.nov – D. tex: P* = 0.093; *D.am – D.tex: P* = 0.840) were not significant. Given the high contribution of PC1 and PC2 to total environmental variation (75.8%, Table S1 and below) and their significant associations with our species, we focused all subsequent analyses of environmental variation on these two axes, and did not further consider PC3.

**Figure 2:**
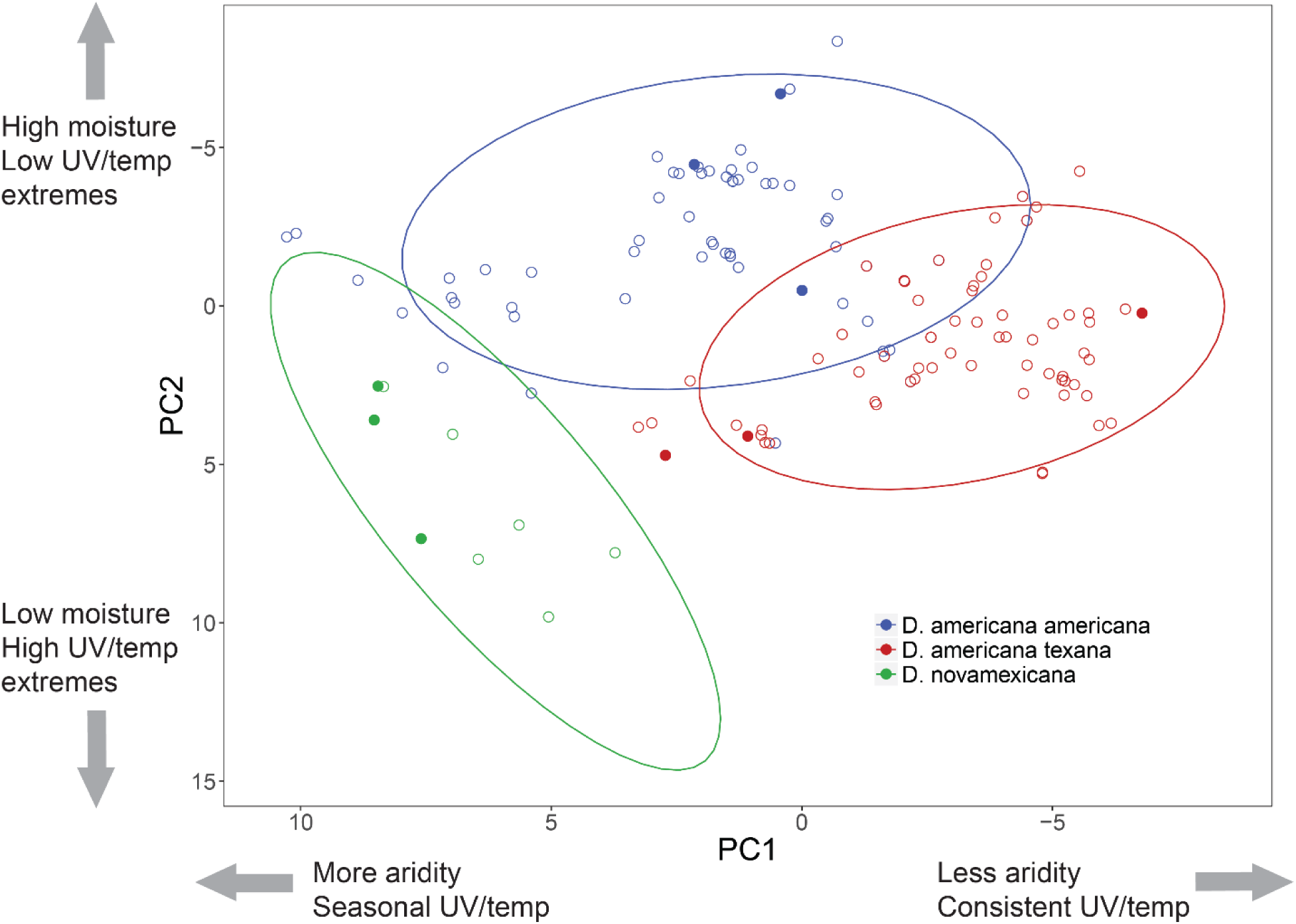
PC1 and PC2 values for all collection records (N=149). PC axes are inverted to mirror spatial orientation of populations in geographic space. Filled circles indicate the 9 focal populations used to assess trait variation. Across all collection records, species differ along PC1 (*F*(2, 82.2, *P* < 0.001) and PC2 (*F*(2, 59.76), *P* < 0.001). Post-hoc Tukey tests indicated that all 3 species differed from one another for both PC1 and PC2 (all *P* values < 0.001).

Based on variable loadings (provided in the supplement; Table S1) we interpret the environmental significance of each of the first 2 PCs as follows: PC1 (explaining 46.0% variance) was strongly negatively loaded for the majority of our precipitation and moisture index variables – with the exception of the seasonality terms; accordingly, higher PC1 values indicate more arid areas with lower rainfall year-round, and higher daily and seasonal temperature variability. PC2 (explaining 29.8% variance) was most strongly loaded for radiation and temperature extremes for the positive values, and negatively loaded for ground moisture variables and temperature/UV seasonality. We therefore interpret high values of PC2 to indicate year-round high temperature and radiation, with low moisture, and negative values to indicate areas with year-round high moisture along with high seasonality in temperature and radiation.

### Traits differed among both species and populations and, in some cases, sexes

Desiccation resistance differed between species (*F*(2, 29.8); *P* < 0.001) as well as between populations within species (*F*(6, 4.5); *P* < 0.001), but not between sexes (*F*(1, 0.8); *P* = 0.364); dry weight (body size) had no effect (*F*(1, 1.3), *P* = 0.25). Post-hoc Tukey tests indicated significant pairwise differences between all three species (*D.nov – D. am: P* < 0.001, *D.nov* – *D. tex*: *P* < 0.001; *D.am* – *D. tex: P* = 0.002); *D. novamexicana* had the greatest desiccation resistance, followed by *D. a. texana* and *D. a. americana* (Figure 3). Consistent with this, *D. novamexicana* populations generally performed better than the other populations, with the exception of one *D. a. texana* population (Morrilton, Arkansas) which had the third highest survival overall (Figure 4).

**Figure 3:**
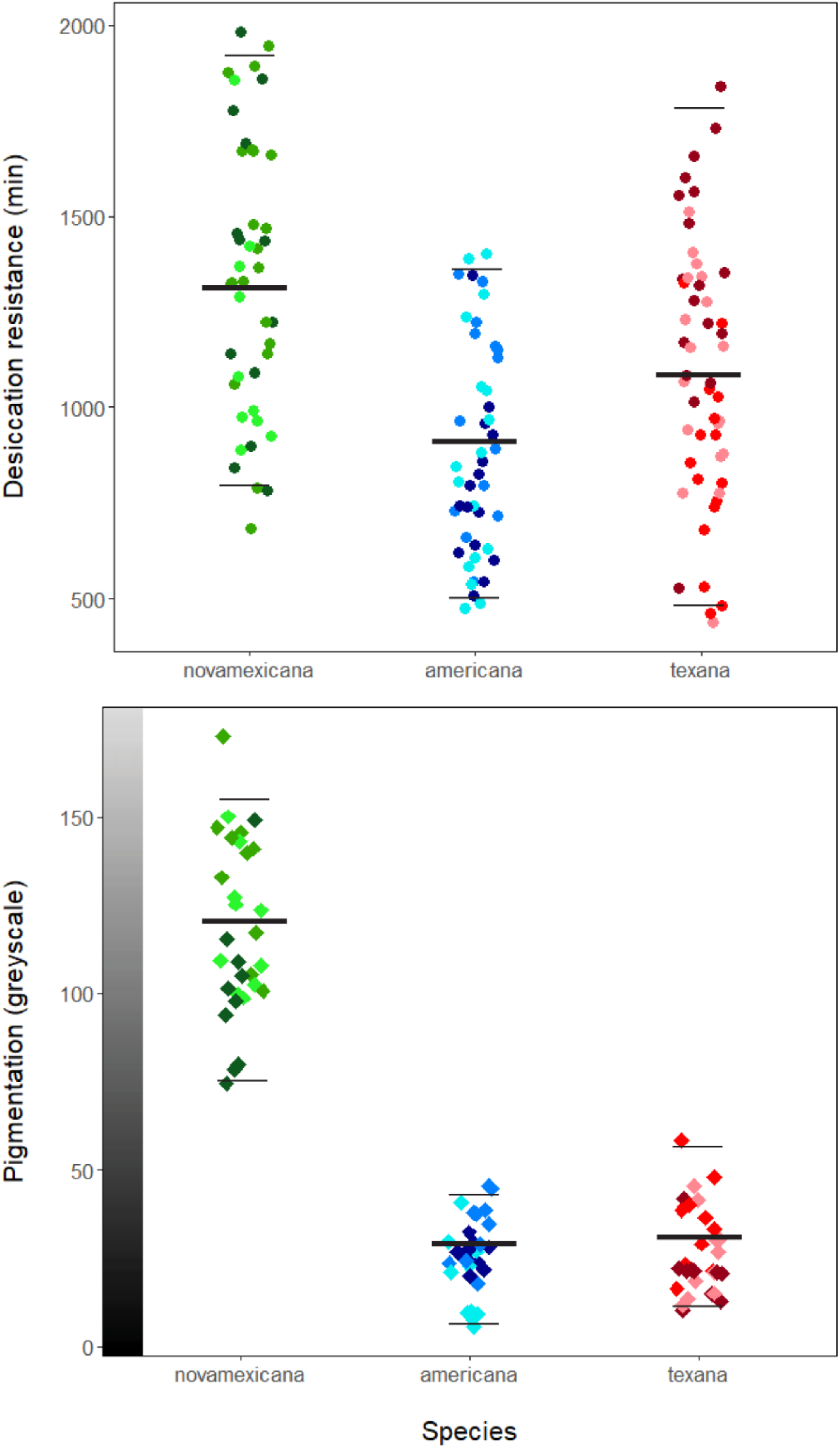
Distribution of individual values for desiccation resistance (top, circles) and pigmentation (bottom, diamonds), according to species (X axis) and population within species (shade variation among points). The y-axis values for individual cuticles on the pigmentation graph correspond to a computed greyscale value represented by the gradient bar. The thick bars indicate species means; thin bars indicate 95% confidence intervals around the mean. Desiccation resistance differs between species (*F*(2, 29.76); *P* < 0.0001) and populations within species (*F*(6, 4.48); P = 0.0004). Post-hoc tests indicate significant differences in all pairwise contrasts (*D.nov* – *D. am*: *P* < 0.0001, *D.nov* – *D. tex*: *P* = 0.00039; *D.am* – *D. tex: P* = 0.0023). Pigmentation differs between both species (*F*(2, 11.86), *P* < 0.0001) and populations within species (*F*(6, 3.13), *P* = 0.0083). Post-hoc contrasts indicate *D. novamexicana* is significantly lighter than *D. a. americana* (*P* < 0.0001) and *D. a. texana* (*P* < 0.0001); *D. a. americana* and *D. a. texana* do not differ (*P* = 0.96).

**Figure 4:**
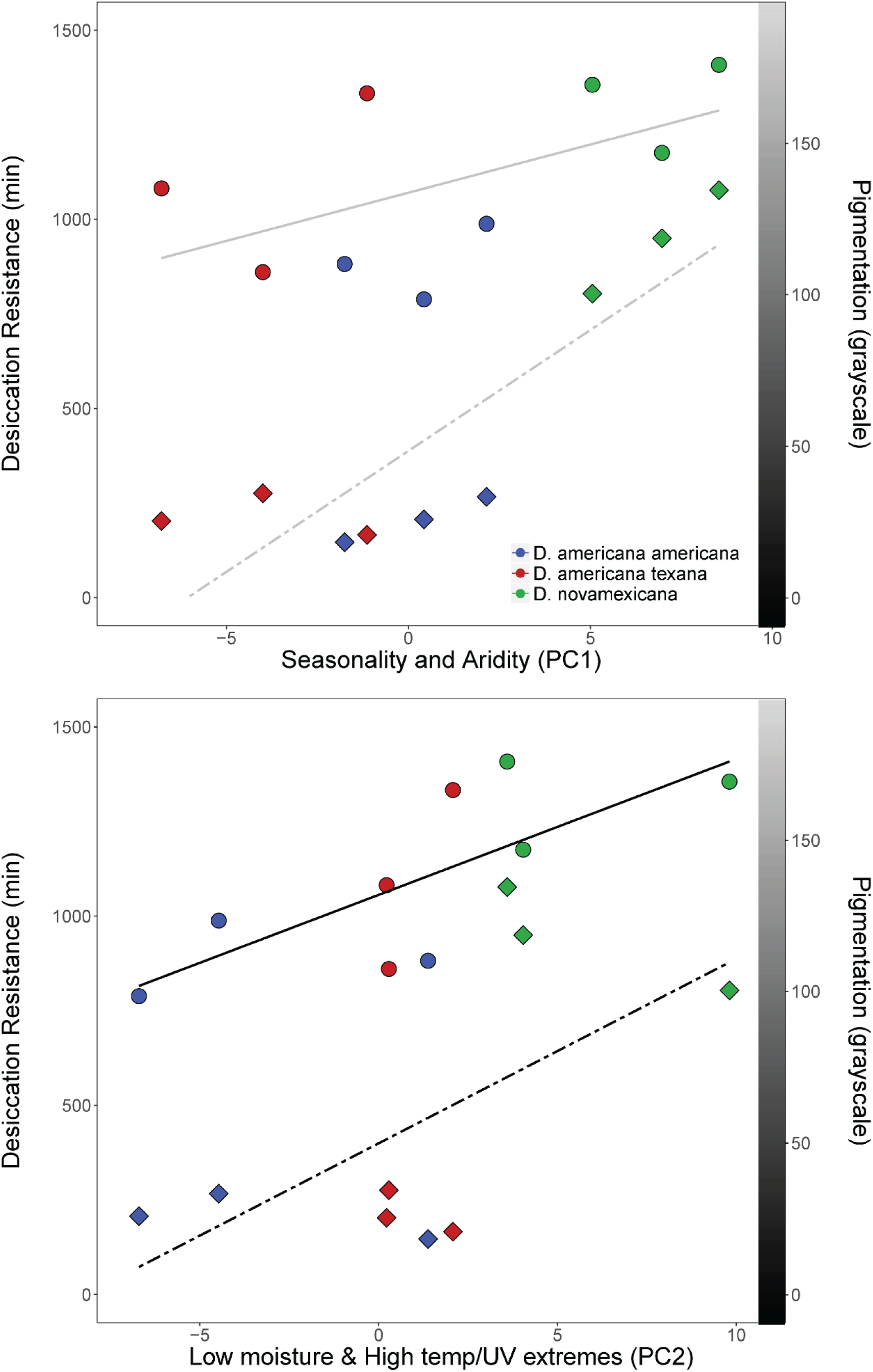
Relationship between mean desiccation resistance (circles) or pigmentation (diamonds) in each population and PC1 (top panel) or PC2 (bottom panel). The left-side y-axis corresponds to desiccation resistance in minutes survived, while the right-side y-axis show pigmentation values on that correspond to a greyscale value represented by the gradient bar. PC1 is not significantly associated with desiccation resistance (r(7) = 0.15; *P* = 0.71) or pigmentation (r(7) = 0.53; *P* = 0.15) was across all 9 populations (top, grey trend lines). PC2 is associated with both desiccation resistance (r(7) = 0.74; *P* = 0.022) and pigmentation (r(7) = 0.68; *P* = 0.044) (bottom, black trend lines), prior to multiple testing correction.

We found that UV resistance was significantly influenced by sex (*F*(1, 4.22), *P* < 0.001) with females living longer in general, and experiencing lower reduction in life expectancy with increasing UV exposure, compared to males. Among energy levels we found that only the 5000J exposure group (*F*(3, 12.36), *P* < 0.001) significantly differed in relative survival from the reference treatment level (100J, see methods), while 500J (*F*(3, −0.23), *P* = 0.82) and 1000J (*F*(3, −0.31), *P* = 0.76) did not. We also found that *D. novamexicana* (*F*(2, −0.363), *P* = 0.72) and *D. a. texana* (*F*(2, 1.41), *P* = 0.16) did not significantly differ from *D. americana*. For full survival curves for each population, sex, and treatment, see Figure S1.

Abdominal pigmentation differed between species (*F*(2, 11.9), *P* < 0.001) and between populations within species (*F*(6, 3.1), *P* = 0.008), and marginally differed between sexes (*F*(1, 0.06), *P* = 0.063). *D. novamexicana* were the lightest flies, and post-hoc Tukey tests confirmed *D. novamexicana* was significantly less pigmented than *D. a. americana* (*P* < 0.001) or *D. a. texana* (*P* < 0.001), while the two americana subspecies had similar pigmentation (*P* = 0.96) (Figure 3). With respect to sex-specific differences, females were marginally less pigmented than males.

### Modest associations between physiological trait variation and major axes of environmental variation

Across all nine populations, we found modest correlations between PC2 and both desiccation resistance (r(7) = 0.74; *P* = 0.022) and pigmentation (r(7) = 0.68; *P* = 0.044), although these do not survive multiple test correction (Figure 4). Neither trait was associated with PC1 (desiccation: r(7) = 0.15; *P* = 0.71; pigmentation: r(7) = 0.53; *P* = 0.15) (Figure 4). For UV, among all associations tested between PCs and median death at each treatment level, including the control 0J exposure treatment (longevity), the only detected correlations were between PC1 and both 100J treatment survival (r(7) = 0.70; *P* = 0.036) and longevity (r(7) = 0.65; *P* = 0.060), though none of the tests survived Bonferroni-correction. All other UV results are provided in the supplement (Table S2).

When taking into account both species and population-level effects on trait variation, for both PCs we found that species differences explained desiccation resistance variation among males but not females (Table 2); in comparison, intraspecific variation in desiccation resistance was not associated with residuals of either PC for either sex (Table 2). For pigmentation variation, in addition to significant differences among species for both sexes, we found that intraspecific variation was modestly associated with PC2 for both males and females, although not with PC1 (Table 2). Based on the factor loadings for PC2 (see methods and Table S1), this latter finding suggests that higher year-round UV exposure and temperature coupled with low moisture (i.e. positive values of PC2), are associated with relatively darker pigmentation within each of these species. Unlike the other two traits, for UV resistance, we found little evidence for consistent associations between quantitative trait variation and macroecological variation, regardless of sex or treatment level. We found only one significant association after correcting for multiple testing: for the 5000J treatment, females showed a significant association with PC2 (F(1) = 12.75; P = 0.009). Complete results of UV analyses are provided in the supplemental material (Table S3).

**Table 1:** Associations between principal component (PC) axes of environmental variation and physiological traits for these species. Reported statistics are from multi-way ANOVA models that include a PC (d.f.=1) and species (d.f.=2) as independent variables, and a given physiological trait as the dependent variable. Bonferroni corrected significance level is *P* < 0.0004.

|  | Desiccation Resistance |  |  |  | Pigmentation |  |  |  |
| --- | --- | --- | --- | --- | --- | --- | --- | --- |
|  | Male |  | Female |  | Male |  | Female |  |
|  | *F* | *P* | *F* | *P* | *F* | *P* | *F* | *P* |
| **PC1** | 0.23 | 0.65 | 0.05 | 0.83 | 0.98 | 0.37 | 0.99 | 0.36 |
| **Species** | 6.29 | 0.043 | 2.46 | 0.18 | 45.04 | 0.00063 | 79.4 | 0.00016 |
| **PC2** | 0.60 | 0.47 | 0.97 | 0.37 | 7.26 | 0.043 | 6.53 | 0.051 |
| **Species** | 6.73 | 0.038 | 2.91 | 0.14 | 92.33 | 0.00011 | 152.6 | <0.0001 |

**Table 2:** Associations between principal component (PC) axes of environmental variation and UV resistance, split by treatment. Reported statistics are from multi-way ANOVA models that include a PC (d.f.=1) and species (d.f.=2) as independent variables, and a given physiological trait as the dependent variable. Bonferroni corrected significance level is *P* < 0.0004.

| Energy | *Females* |  |  |  |  |  |  |  |  |  |
| --- | --- | --- | --- | --- | --- | --- | --- | --- | --- | --- |
|  | 0J |  | 100J |  | 500J |  | 1000J |  | 5000J |  |
|  | *F* | *P* | *F* | *P* | *F* | *P* | *F* | *P* | *F* | *P* |
| **PC1** | 4.83 | 0.079 | 3.62 | 0.12 | 0.75 | 0.43 | 0.33 | 0.59 | 0.076 | 0.79 |
| **Species** | 1.91 | 0.24 | 3.16 | 0.13 | 1.51 | 0.31 | 1.21 | 0.37 | 2.57 | 0.17 |
| **PC2** | 6.45 | 0.052 | 4.09 | 0.10 | 2.93 | 0.15 | 0.004 | 0.95 | 3.18 | 0.13 |
| **Species** | 2.22 | 0.20 | 3.33 | 0.12 | 2.09 | 0.22 | 1.14 | 0.39 | 4.14 | 0.09 |
| Energy | *Males* |  |  |  |  |  |  |  |  |  |
|  | 0J |  | 100J |  | 500J |  | 1000J |  | 5000J |  |
|  | *F* | *P* | *F* | *P* | *F* | *P* | *F* | *P* | *F* | *P* |
| **PC1** | 0.94 | 0.38 | 0.76 | 0.42 | 3.73 | 0.11 | 3.20 | 0.13 | 0.70 | 0.70 |
| **Species** | 0.33 | 0.73 | 2.69 | 0.16 | 7.59 | 0.03 | 0.39 | 0.70 | 0.18 | 0.18 |
| **PC2** | 1.48 | 0.28 | 5.31 | 0.069 | 1.04 | 0.35 | 1.78 | 0.24 | 0.045 | 0.045 |
| **Species** | 0.36 | 0.71 | 4.81 | 0.068 | 5.25 | 0.059 | 0.32 | 0.74 | 0.052 | 0.052 |

For associations between traits, we found that higher desiccation resistance is positively correlated with lighter pigmentation in males (r(7) = 0.70, *P* = 0.036), but only marginally for females (r(7) = 0.59, *P* = 0.091)(Figure S2). In contrast, UV resistance was not correlated with pigmentation for either sex at any of the treatment levels (Table S4). UV resistance was modestly negatively correlated with desiccation resistance for females at the highest exposure (5000J, r(7) = −0.720, *P* = 0.029), but this association does not survive multiple-testing correction.

## Discussion

Here we examined differences in environment between occurrence locations of three closely related North American *Drosophila* species and then evaluated three physiological traits that might have been shaped by these climatic differences. We found that all three species differed along two major principal axes of environmental variation that were strongly loaded for precipitation and ground moisture variables. We also detected differences between species in desiccation resistance and abdominal pigmentation, but not in assayed UV resistance. Although there was little evidence for finer-scale trait-environment associations within species, broader species differences—especially in desiccation resistance between *D. novamexicana* and its relatives—are consistent with differential natural selection imposed by habitats that vary in their frequency of rain and consistency of moisture. Quantifying macroecological variation across the ranges of these closely related species, as well as the accompanying patterns of trait variation, therefore enabled us to identify potentially important agents of climate selection and the scales at which they might be shaping adaptive physiological differences.

### Water availability characterizes spatial variation and habitat divergence in species distributions

Identifying environmental factors that differentiate closely related species is essential for understanding the selective agents that could act on trait variation between, and possibly within, these species. Here we confirmed that *D. novamexicana* is environmentally differentiated from both *D. americana* and *D. texana*, consistent with strong habitat differences between this largely desert-associated species and its non-xeric relatives. Along our largest axis of environmental variation (PC1, that explained 46.0% of the variance in bioclimatic variables) *D. novamexicana* had the highest PC1 values, indicative of habitats with low year-round rainfall and ground moisture, and higher daily and annual temperature and UV fluctuations. Interestingly, this same major axis also differentiated the southern *D. a. texana* from the northern *D. a. americana* subspecies; *D. a. texana* had the lowest values of the three species, with population locations characterized by consistently heavier precipitation and moisture, and smaller temperature fluctuations (Figure 2). Our finding that the same axis differentiates our two *D. americana* subspecies points to the importance of water availability and covarying temperature/UV factors in defining geographic differences among all three taxa in this group. This inference is further supported by similarly strong differentiation between each species in the second largest axis of environmental variation, PC2. High values of PC2 are associated with high peak temperature and UV values coupled with low ground moisture, and so can be interpreted as dry heat and heavy sun exposure versus humid warm periods with more consistent and lower peak UV intensity. While this axis is not loaded strongly for precipitation *per se*, moisture still stands out as explanatory.

These data combined indicate the broad importance of water availability for delineating the geographic distribution of all three species in this group, beyond simple spatial separation. The most strongly differentiating bioclimatic factors—variation in precipitation and moisture, but also in temperature and UV—are strong candidates as broad selective agents that might shape trait variation among these species. Because these macroecological factors themselves generate expectations about the kinds of traits that might be responding to them—namely, physiological traits associated with adaptive responses to moisture, UV, and temperature variation—we directly evaluated these expectations with relevant trait data within and between our species.

### Desiccation resistance and pigmentation vary with macroecological variation, but patterns differ between traits

Given the macroecological differences between our species, we assayed variation in three traits to evaluate evidence for climate selection shaping adaptive trait variation: desiccation resistance, UV resistance, and pigmentation. All three have previously been proposed as targets of natural selection imposed by environmental variation (Matute and Harris 2013, Wittkopp et al. 2011, Karan and Parkash 1998, Bastide et al. 2014, Parkash et al. 2008), either within or between closely related *Drosophila* species. Here we similarly found evidence that variation across species in desiccation resistance accompanies macroecological differences in moisture availability. Across our nine focal populations, mean survival times under acute desiccation were associated with our PC2 environmental axis (Figure 4); that is, populations that experience year-round high temperature and UV exposure with little ground moisture (high PC2 values) have consistently higher desiccation resistance than those from regions with more consistent moisture availability. Moreover, we infer that this association is largely driven by differences in habitat-imposed selection among species. In particular, *D. novamexicana* was characterized by the most xeric habitats and had significantly elevated desiccation resistance relative to its mesic sister species, presumably because this confers improved survivorship throughout a substantially drier species range. More generally, significant species differentiation both in trait and environment, and a trait-environment correlation across populations, further strongly suggest that moisture and other climatic factors associated with PC2 are causal selective agents acting on desiccation resistance variation among all three species examined.

In contrast to desiccation resistance, no consistent trait-environment association emerged from our analysis of UV resistance, nor were species consistently different, although we did detect a significant difference between sexes. This lack of association with macroecological variation is interesting because, just as for moisture and precipitation, our two major PC axes are also loaded for UV intensity variables. This result might reflect the limitations of extrapolating natural functional differences in UV tolerance from survivorship after an acute single exposure to UV (a technique that is commonly used; Matute and Harris 2013, Jacobs 1974, Wang et al. 2008). A more realistic assay, involving periodic or sustained exposures of lower average dose, might better simulate natural differences in daily UV exposure associated with macroecological conditions. Nonetheless, despite its potential shortcomings, we note that our assay did reveal differences in both relative and absolute survival of males compared to females; across all species, females consistently show a less drastic reduction in lifespan relative to males, even at the highest UV exposures (Figure S3). This suggests that our assay was sufficient to capture some aspects of sex-specific biological variation in physiological responses to UV, and indicates that females in this group show greater high-dosage UV resistance than males. Finally, in addition to greater UV resistance in females, we also detected greater female longevity in the absence of UV exposure (i.e. at the control, 0J exposure). Greater female longevity is a pattern seen across many, but not all, *Drosophila* species (Yoon et al. 1990, Durbin and Yoon 1986) and several factors--including sex-linkage of the underlying causal factors--could explain sex differences in baseline lifespan, beyond life history differences associated with ecological variation.

Our analysis of pigmentation variation revealed yet a third pattern of trait variation among our populations and species. While we confirmed that differences in abdominal pigmentation in this group can be stark—*D. novamexicana* has a light almost tan appearance that is quantitatively different from the dark brown to black of the two *D. americana* subspecies—we also found that pigmentation intensity did not differ between the *D. americana* and *D. texana* populations we examined here, unlike differences detected in desiccation resistance. Moreover, while mean pigmentation intensity was associated with environmental axis PC2 (*P* = 0.044) across all nine populations, this pattern of association was effectively bimodal: *D. novamexicana* populations had high values for PC2 and low pigmentation intensity, while the lower PC2 values for populations of both other species were accompanied by high, relatively invariant, pigmentation intensity (Figure 3, 4). Pigmentation also differed from desiccation resistance in that we found evidence for sex differences in this trait, that were absent for desiccation resistance. These results indicate that, although there is evidence that pigmentation covaries with macroecological climate variation (especially PC2), its specific association differs from desiccation resistance. This could be due to a number of factors, including the genetic architecture of pigmentation itself. Wittkopp et al. (2009) have shown that the yellow body color of *D. novamexicana* is a fixed, derived phenotype determined by alleles at the *tan* and *ebony* loci, and inferred that these alleles pre-date speciation, as they can also be found segregating across populations of *D. americana.* Furthermore, introgressing either or both of these loci from *D. americana* into *D. novamexicana* produced darker morphs of *D. novamexicana,* with pigmentation closer to that of the donor species (Wittkopp et al. 2011). The observation that exchanging just these two alleles can substantially alter pigment phenotypes suggests that the genetic architecture of pigmentation variation might be more simple than desiccation resistance, and less likely to generate phenotypes that vary incrementally with environmental variation. This difference in genetic architecture might contribute to differences in the specific trait-environment associations we observed for pigmentation versus desiccation resistance, even though both traits clearly covary along similar macroecological axes.

Finally, we also note that the covariation between pigmentation and desiccation resistance is most likely shaped by separate responses of each trait to climate variation, rather than by a direct mechanistic connection between them, as has been previously suggested for other *Drosophila* species (Brisson et al. 2005, Rajpurohit et al. 2008, Karan and Parkash 1998, Parkash et al. 2008). Although we did detect a modest correlation between pigmentation and desiccation resistance variation (*P* = 0.036), previous work in this system provides strong evidence that these traits are not mechanistically associated. In particular, in their genetic analysis, Wittkopp et al. (2011) found that introgression of *ebony* or *tan* alleles from *D. americana* into the background of *D. novamexicana* did not alter desiccation resistance. Based on these data, Wittkopp et al. (2011) concluded that relative humidity was unlikely to be the selective factor driving evolution of pigmentation variation in *D. americana*. Clusella-Trullas and Treblanche (2011) reanalyzed this dataset and inferred that mean diurnal temperature range and solar radiation represented the best model for explaining underlying pigmentation variation. From this, they inferred that *D. americana* pigmentation variation might be driven by spatial variation in thermal stress, consistent with the ‘thermal melanism hypothesis’—which proposes that darker pigmentation in colder regions allows ectotherms to increase and maintain body temperature more rapidly. While we did not have as broad a sample of pigmentation variation for *D. americana*, our results indicate some support for this hypothesis: annual solar radiation and mean diurnal range are both loaded highly for PC2, however several other variables not tested by Clusella-Trullas and Treblanche (2011)—most notably ground moisture variables—were also strongly loaded. It therefore seems reasonable that thermal regulation may have played a role in evolution of pigmentation in this system, but that it is likely not the sole factor shaping evolution of this trait.

### Intraspecific trait variation differs from species-level patterns

While we detected differences between species in environmental factors and in two of our traits, our data also clearly revealed substantial trait variation between populations within species, and within populations in some cases. Similar to differences between species, intraspecific variation can reveal signals of local adaptation when environmental variation among populations is associated with relevant trait variation between these populations. However, despite ample intraspecific variation for all three traits, we found little evidence for strong trait-environmental associations at the subspecific level, at least with the limited sample of populations in which we examined these relationships. Indeed, once species differences are accounted for, only pigmentation intensity showed evidence for environmental correlations within species. These associations were modest and, interestingly, in the opposite direction from the pattern detected between species, such that populations with higher PC2 values tended to be more darkly pigmented (Figure 4, Table 2). This curious observation implies that factors that are shaping local patterns of pigmentation might run contrary to the processes that yielded the between-species pattern—a hypothesis that could be evaluated with future work across a broader set of populations.

Apart from limited power, this general lack of strong intraspecific trait-environment associations could be due to several non-exclusive factors, although in the absence of more direct data they remain hypotheses. First, while bioclimatic variables can be very effective in describing broad environmental variation, they do not provide a complete view of local differences in abiotic and biotic environments, although these might nonetheless be critical for shaping trait-environment relationships. Similarly, populations within our species might be responding to local selective conditions via strategies or behaviors that are not observable in the experimental assays we used here. For example, factors like shade availability, proximity to local sources of water, and/or strategies to seek out these areas, may relax or exacerbate local selection on desiccation stress responses in ways that do not match broad macroecological climate variation. A similar proposal was suggested by Sillero et al. (2014) to explain a lack of association between source environment and locomotor activity under thermal stress in these same species. Second, there could be additional within-species constraints on the mechanisms underpinning these traits, such that they are either unable to respond more finely to local conditions, or the fitness benefits of doing so are insufficient to outweigh the benefits of optimizing alternative functions. One example potentially relevant to desiccation resistance variation involves cuticular hydrocarbon (CHC) composition—the blend of waxy compounds on the surface of insects—that is known to be important for desiccation resistance in multiple species, but can also play a role in sexual communication (reviewed Chung and Carroll 2015). If maintaining effective CHC-mediated sexual signaling species-wide imposes constraints on local physiological responses in CHC-mediated desiccation resistance, this could produce a pattern in which species strongly differ in desiccation resistance (in a way that matches broad environmental variation), but there is no strong local pattern of adaptation in the same physiological response. Finally, regardless of the specific selective factors responsible, changes in strength or nature of selection over the evolutionary history of divergence among our species could also cause patterns of variation to appear more complex.

## Conclusions

Our study took a multi-faceted approach to assessing the environmental factors that could shape natural variation in several ecologically-relevant physiological traits. We used broad patterns of bioclimatic variation to generate predictions about physiological trait adaptation between species, that we then evaluated with relevant trait data within and between our species. Our observation that macroecological variation—in particular water availability—covaries with both pigmentation and desiccation resistance indicates that abiotic environmental variation is likely important in the adaptive history of both these traits. Nonetheless, specific patterns of trait-environment association could be influenced by other factors, such as differences in genetic architecture among traits or local variation in ecological factors, highlighting the importance of evaluating these relationships at multiple evolutionary scales.

## Abbreviations

Not applicable

## Declarations

### Ethics approval and consent

Not applicable

### Consent for Publication

Not applicable

### Availability of data and materials

All data generated or analyzed during this study will be uploaded and linked upon acceptance of article.

### Author’s contributions

JSD and LCM conceived and designed the experiment. JSD performed the experiment and statistical analyses. JSD and LCM wrote and edited the manuscript. All authors read and approved manuscript.

## Acknowledgements

We thank the Drosophila species stock center for providing the population lines used in this study. Research was supported by Indiana University Department of Biology funding to LCM and JSD.

