## Supplementary material for "Physiological variation reflects bioclimatic differences in the *Drosophila americana* species complex"

**Table S1: Bioclimatic variables used in analysis of environmental variation within and between species, including source, resolution, and loadings for the first 3 principal component axes.**

|  | Variable | Source | Resolution | PC1 loading | PC2 loading | PC3 loading |
| --- | --- | --- | --- | --- | --- | --- |
| BIO1 | Annual Mean Temp | Worldclim | 30” | -0.163 | 0.218 | -0.046 |
| BIO2 | Mean Diurnal Range | Worldclim | 30” | 0.115 | 0.159 | 0.106 |
| BIO3 | Isothermality | Worldclim | 30” | -0.121 | 0.215 | -0.065 |
| BIO4 | Temperature Seasonality | Worldclim | 30” | 0.188 | -0.171 | 0.085 |
| BIO5 | Max Temperature of Warmest Month | Worldclim | 30” | -0.017 | 0.268 | 0.065 |
| BIO6 | Min Temperature of Coldest Month | Worldclim | 30” | -0.188 | 0.189 | -0.058 |
| BIO7 | Temperature Annual Range | Worldclim | 30” | 0.219 | -0.106 | 0.099 |
| BIO8 | Mean Temperature of Wettest Quarter | Worldclim | 30” | 0.002 | 0.116 | -0.401 |
| BIO9 | Mean Temperature of Driest Quarter | Worldclim | 30” | -0.174 | 0.162 | 0.216 |
| BIO10 | Mean Temperature of Warmest Quarter | Worldclim | 30” | -0.111 | 0.227 | 0.004 |
| BIO11 | Mean Temperature of Coldest Quarter | Worldclim | 30” | -0.174 | 0.213 | -0.057 |
| BIO12 | Annual Precipitation | Worldclim | 30” | -0.237 | -0.069 | 0.059 |
| BIO13 | Precipitation of Wettest month | Worldclim | 30” | -0.221 | 0.020 | -0.107 |
| BIO14 | Precipitation of Driest month | Worldclim | 30” | -0.211 | -0.104 | 0.170 |
| BIO15 | Precipitation Seasonality | Worldclim | 30” | 0.130 | 0.155 | -0.250 |
| BIO16 | Precipitation of Wettest Quarter | Worldclim | 30” | -0.224 | -0.002 | -0.124 |
| BIO17 | Precipitation of Driest Quarter | Worldclim | 30” | -0.215 | -0.098 | 0.175 |
| BIO18 | Precipitation of Warmest Quarter | Worldclim | 30” | -0.199 | -0.035 | -0.276 |
| BIO19 | Precipitation of Coldest Quarter | Worldclim | 30” | -0.217 | -0.047 | 0.222 |
| BIO20 | Annual mean radiation | Climond | 10’ | -0.052 | 0.290 | 0.062 |
| BIO21 | Highest weekly radiation | Climond | 10’ | 0.146 | 0.196 | 0.150 |
| BIO22 | Lowest weekly radiation | Climond | 10’ | -0.121 | 0.263 | -0.028 |
| BIO23 | Radiation seasonality | Climond | 10’ | 0.173 | -0.208 | 0.050 |
| BIO24 | Radiation of wettest quarter | Climond | 10’ | 0.114 | 0.035 | -0.348 |
| BIO25 | Radiation of driest quarter | Climond | 10’ | -0.103 | 0.165 | 0.320 |
| BIO26 | Radiation of warmest quarter | Climond | 10’ | 0.156 | 0.172 | 0.180 |
| BIO27 | Radiation of coldest quarter | Climond | 10’ | -0.130 | 0.254 | -0.047 |
| BIO28 | Annual mean moisture index | Climond | 10’ | -0.204 | -0.165 | 0.028 |
| BIO29 | Highest weekly moisture index | Climond | 10’ | -0.208 | -0.139 | 0.064 |
| BIO30 | Lowest weekly moisture index | Climond | 10’ | -0.171 | -0.196 | -0.073 |
| BIO31 | Moisture index seasonality | Climond | 10’ | 0.035 | 0.137 | 0.263 |
| BIO32 | Mean moisture index of wettest quarter | Climond | 10’ | -0.208 | -0.142 | 0.074 |
| BIO33 | Mean moisture index of driests quarter | Climond | 10’ | -0.191 | -0.171 | -0.102 |
| BIO34 | Mean moisture indeox of warmest quarter | Climond | 10’ | -0.200 | -0.092 | -0.241 |
| BIO35 | Mean moisture index of coldest quarter | Climond | 10’ | -0.188 | -0.154 | 0.169 |

**Table S2: Associations between environmental PC1 or PC2 and UV survival, separated by treatment level. Reported statistics are from Pearson’s correlation between each PC and median UV resistance. Bonferroni-corrected significance level is *P* < 0.0004.**

| **Energy** | **PC1** | | **PC2** | |
| --- | --- | --- | --- | --- |
|  | *r* | *P* | *r* | *P* |
| 0J | 0.48 | 0.20 | 0.37 | 0.42 |
| 100J | 0.64 | 0.061 | -0.037 | 0.92 |
| 500J | 0.10 | 0.79 | -0.0011 | 0.99 |
| 1000J | 0.53 | 0.14 | 0.12 | 0.76 |
| 5000J | 0.092 | 0.81 | -0.0003 | 0.99 |

**Table S3: Correlations between UV resistance (survival in days, post-exposure) at each energy exposure level and each of the other two physiological traits measured. Reported statistics are Pearson’s correlation coefficients between each trait and UV survival at one energy level, separated by sex. Bonferroni-corrected significance level is *P* < 0.0004.**

| **Energy** | **Pigmentation** | | | | **Desiccation Resistance** | | | |
| --- | --- | --- | --- | --- | --- | --- | --- | --- |
|  | **Males** | | **Females** | | **Males** | | **Females** | |
|  | r | *P* | r | *P* | r | *P* | r | *P* |
| 0J | -0.063 | 0.872 | -0.652 | 0.057 | -0.079 | 0.84 | -0.226 | 0.558 |
| 100J | -0.029 | 0.941 | -0.439 | 0.238 | 0.161 | 0.700 | -0.398 | 0.289 |
| 500J | -0.494 | 0.177 | -0.526 | 0.146 | -0.298 | 0.437 | -0.528 | 0.144 |
| 1000J | -0.220 | 0.571 | -0.068 | 0.862 | 0.370 | 0.327 | -0.271 | 0.481 |
| 5000J | -0.394 | 0.295 | 0.704 | 0.034 | -0.555 | 0.121 | 0.563 | 0.115 |

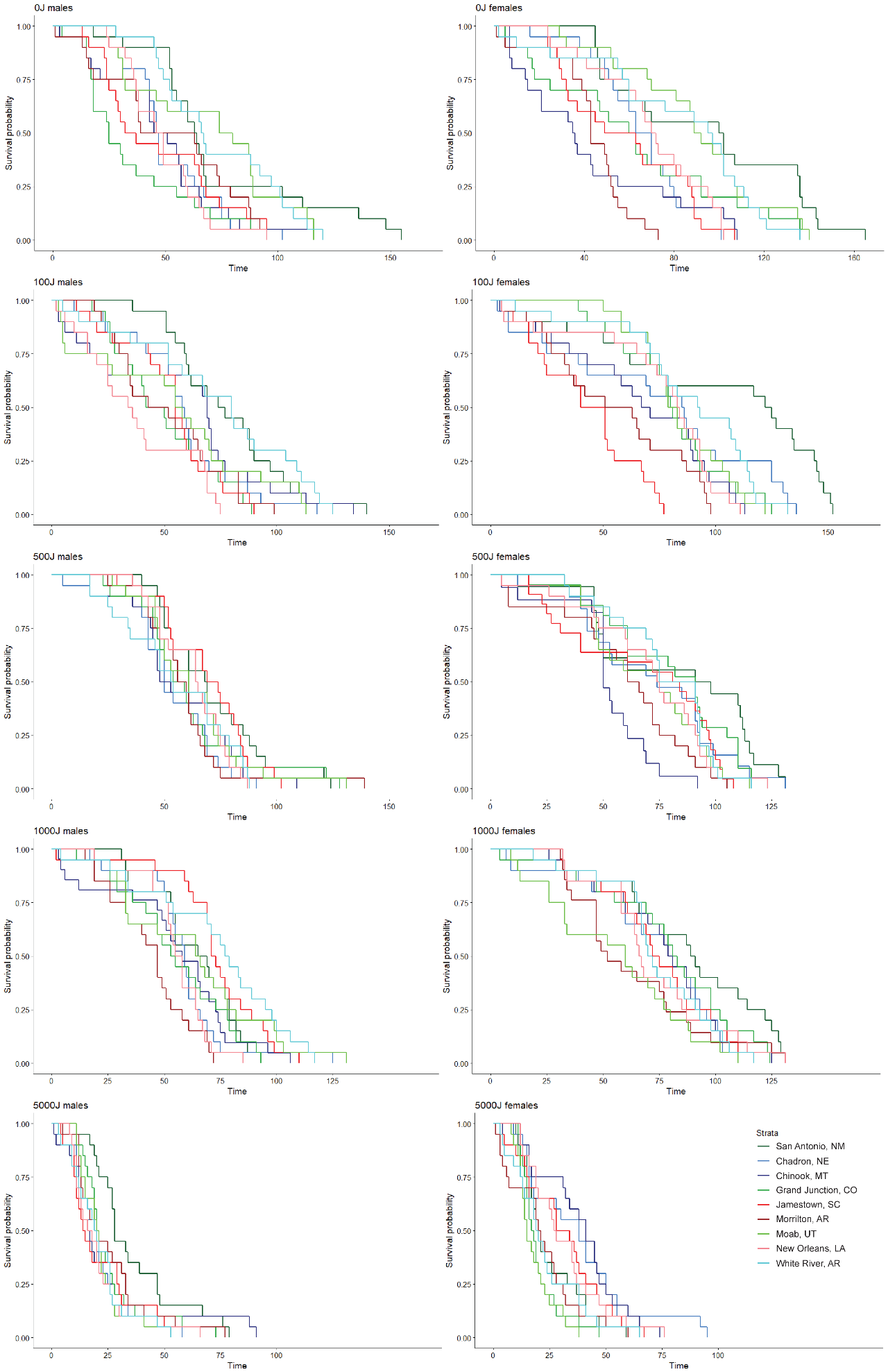

**Figure S1: Meier-Kaplan survival curves after UV treatment for each population. X-axis is time (in days); Y-axis is survival probability (for each 20 fly trial). Panels are organized by UV-B energy exposure (from top to bottom: 0J, 100J, 500J, 1000J, and 5000J) and by sex (with males on the left and females on the right). Population legend is in bottom right for reference throughout.**

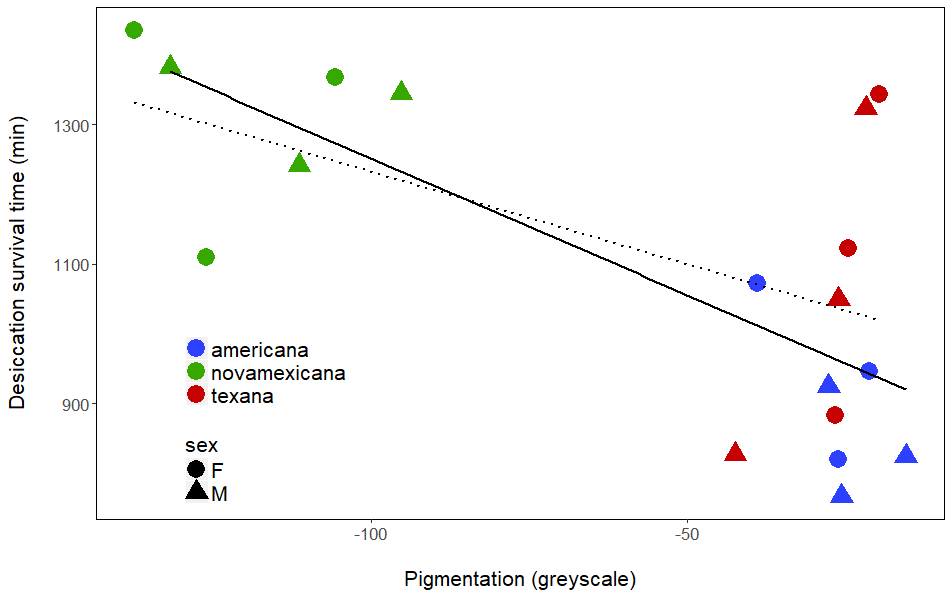

**Figure S2: Associations between desiccation resistance and pigmentation for males (triangles, solid line) and females (circles, dotted line). Mean population desiccation resistance and pigmentation are correlated in males (r(7) = 0.701, *P* = 0.0355), but only marginally in females (r(7) = 0.594, *P* = 0.0914).**

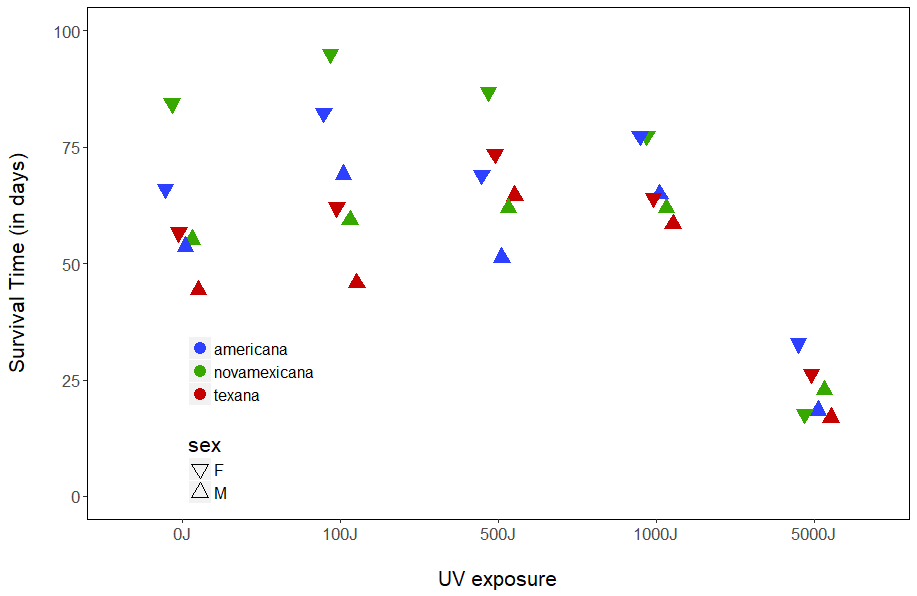

**Figure S3: Survival time in days for UV resistance trials across treatments for each species – split by sex. Survival time was significantly influenced by sex (*F*(1, 4.22), *P* < 0.001) with females living longer in general. With sexes combined, only the 5000J exposure group (*F*(3, 12.36), *P* < 0.001) significantly differed in relative survival from the reference treatment level (100J, see methods), while 500J (*F*(3, -0.23), *P* = 0.82) and 1000J (*F*(3, -0.31), *P* = 0.76) did not.**
